# Neglecting normalization impact in semi-synthetic RNA-seq data simulation generates artificial false positives

**DOI:** 10.1101/2022.05.10.490529

**Authors:** Boris P Hejblum, Kalidou Ba, Rodolphe Thiébaut, Denis Agniel

## Abstract

By reproducing differential expression analysis simulation results presented by Li *et al*, we identified a caveat in the data generation process. Data not truly generated under the null hypothesis led to incorrect comparisons of benchmark methods. We provide corrected simulation results that demonstrate the good performance of dearseq and argue against the superiority of the Wilcoxon rank-sum test as suggested by Li *et al*. Please see related Research article with DOI 10.1186/s13059-022-02648-4.

---

Li *et al* [1] recently raised significant concerns regarding popular RNA-seq differential expression methods `edgeR`[2] and `DESeq2`[3] in the context of large human population sample sizes. We share those concerns, having ourselves come to similar conclusions[4] before, as have others[5, 6]. However, their findings that other methods (namely `dearseq`, `limma-voom`[7], and NOISeq[8]) also have increased false positive rates does not appear to be correct, and the evidence does not support their claim that the Wilcoxon rank-sum test should be preferred to these alternatives. We used the same semi-synthetic datasets that were used in Li *et al* to show that no methods (including Wilcoxon test) are able to maintain the nominal level of “false discoveries” according to their definition because the data used for analysis are not truly generated under *H*_0_. We demonstrate how their permutation scheme should be amended to support analysis of false positive rates under *H*_0_. Using this amended scheme, we show that dearseq appears to outperform other methods.

First, we demonstrate that Wilcoxon test has the same properties as the competing methods when given the same data. We recreated Figure 2A from Li *et al* where the empirical (“actual”) False Discovery Rate (FDR) is plotted against the nominal (“claimed”) FDR using semi-synthetic data generated from the full *GTEx Heart atrial appendage* (n=372) *VS Heart left ventricle* (n=386) simulation[1] in our Figure 1A. We recomputed those results using code and data shared by the authors[9]. The key difference is that we applied the Wilcoxon test on the same normalized data (following the edgeR pipeline for filtering out genes with low counts and using log2-counts per million transformation) used by all other methods. When given the same data as all other methods, the Wilcoxon test also appeared to exaggerate the FDR, as did all other methods.

**Figure 1:**
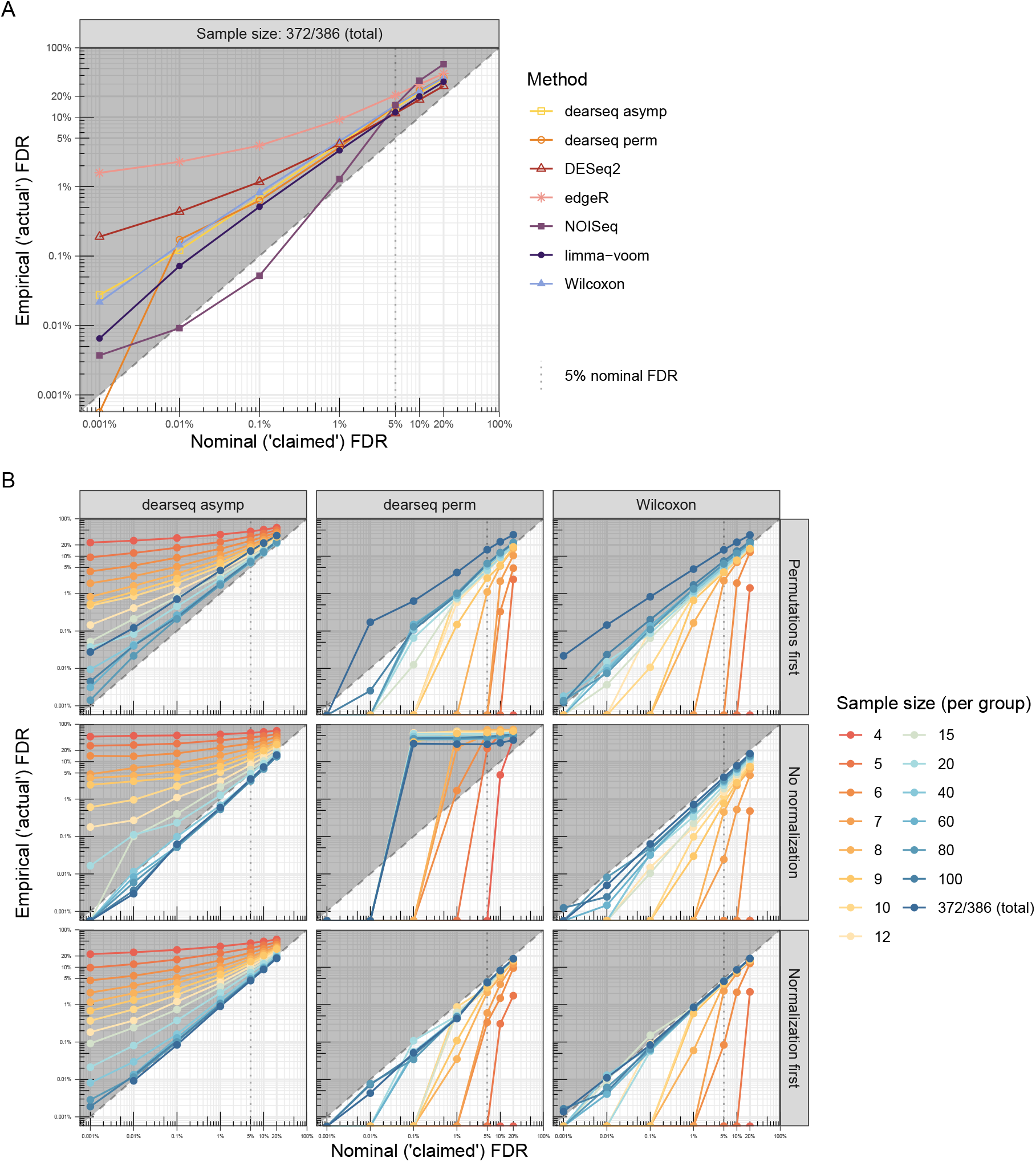
Empirical FDR control against Nominal FDR level. Average over 50 semi-synthetic dataset generated from the *GTEx Heart atrial appendage VS Heart left ventricle* data. 50% of the true Differentially Expressed (DE) genes are randomly sampled in each semi-synthetic dataset (i.e.2,889 genes remains unpermuted as true positives) and considered as gold-standard DE genes. **Panel A** reproduces the results from Li *et al* [1] Figure 2A when all methods are applied to the same data (first permuted to generate null gene expression and then normalized) on the full sample size (372 and 386 samples in each group respectively). **Panel B** study the impact of both the sample size as well as the respective order between the data normalization and the random permutations to generate non-differentially expressed genes on the FDR control of the Wilcoxon test and on both asymptotic and permutation tests from `dearseq`. Of note, only the asymptotic test from `dearseq` works on non-normalized data, while its permutation test fails because of this same issue: observations across samples are not exchangeable and therefore the null distribution is not well estimated by permutations.

**Figure 2:**
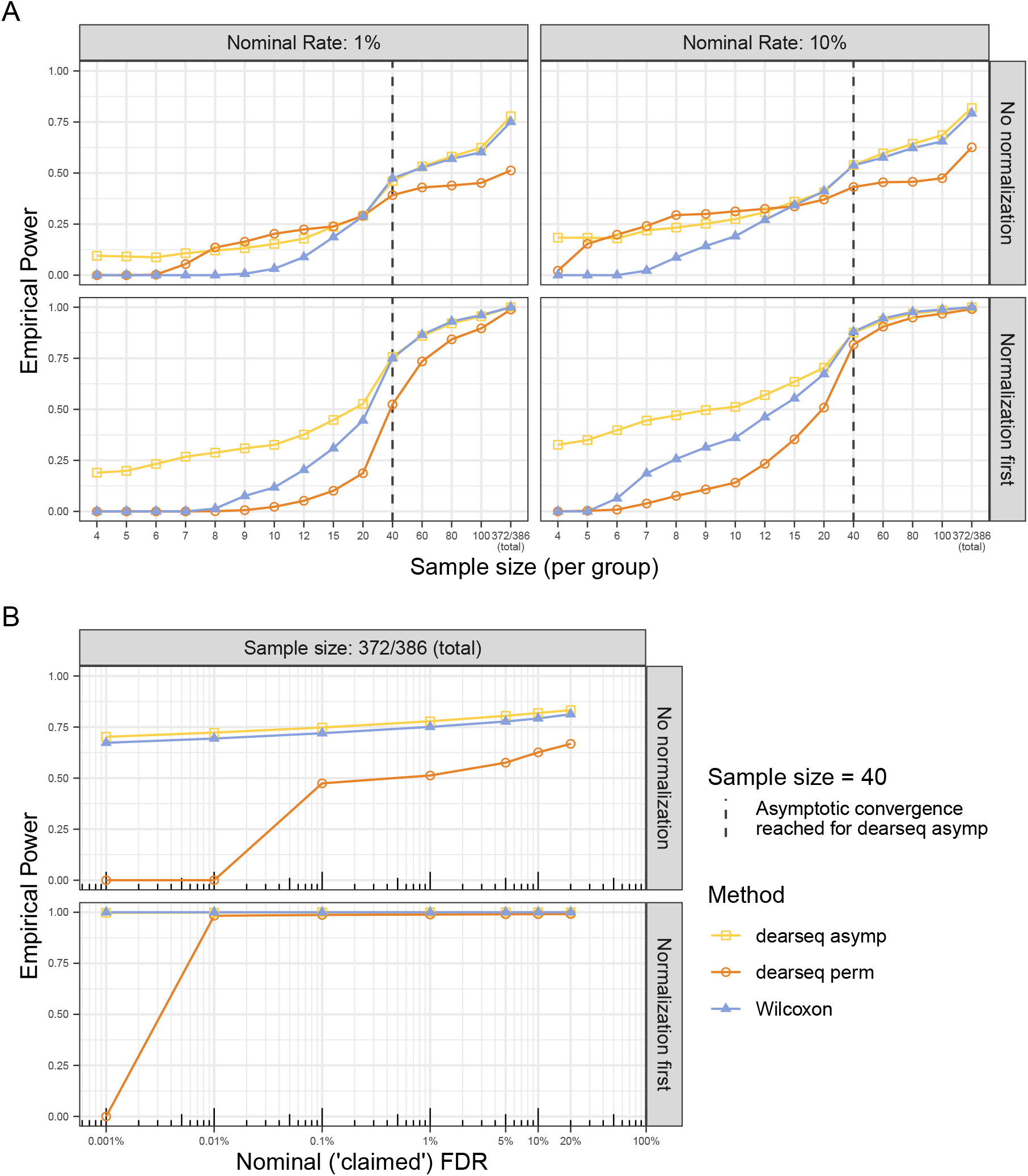
Empirical statistical power for the Wilcoxon test and dearseq asymptotic and permutation tests. Average over 50 semi-synthetic dataset generated from the *GTEx Heart atrial appendage VS Heart left ventricle* data. 50% of the true Differentially Expressed (DE) genes are randomly sampled in each semi-synthetic dataset (i.e. 2,889 genes remains unpermuted as true positives) and considered as gold-standard DE genes used as true positives. **Panel A** reproduces the results from Li *et al* [1] Figure 2B as a function of sample size for both 1% and 10% nominal FDR levels, when all three methods are applied to the same data (either without any normalization or when the data are first normalized before randomly swapping values to generate expressions under *H*_0_ – i.e. the two cases for which the FDR is controlled and thus the empirical power is interpretable). **Panel B** study the impact of the nominal FDR level in both cases for the full sample size (372 and 386 samples in each group respectively).

This apparent increase in FDR was not due to the methods but rather to an inappropriate datageneration scheme. In Figure 1B, we compare the performance of both dearseq asymptotic and permutation tests with the Wilcoxon test across various sample sizes (in their discussion, Li *et al* [1] advocate for permutation analysis, fortunately dearseq already features such a permutation approach which we added to the comparison). In these semi-synthetic data sets, gene expression under *H*_0_ was generated by randomly swapping expression values between samples. However, Li *et al* did not analyze these data directly, but instead normalized them before analysis. The top panel of Figure 1B shows how the Li *et al* permutation scheme leads to an apparent increase in FDR because the expression is no longer generated from *H*_0_ after normalization (e.g. due to a high count being swapped into a sample with a much lower library size, artificially creating a large expression post-normalization). When the data were analyzed without normalization – an approach that would never be used in practice – we show in the middle panel of Figure 1B that both `dearseq` and the Wilcoxon test attained the nominal FDR as sample size increased.

We also show in Figure 1B bottom panel an alternative permutation scheme which fixes the issues with the scheme in Li *et al*: when counts are first normalized before being permuted under *H*_0_, we demonstrate that all three tests adequately controlled the FDR for the full dataset. Figure 2 shows that once convergence was reached, the `dearseq` asymptotic test achieved slightly higher statistical power than Wilcoxon test, while the Wilcoxon test had superior power to `dearseq` permutation test. See Supplementary Figure S3 for statistical power against empirical FDR. This amended permutation scheme should be preferred to the Li *et al* permutation scheme. It is fundamental to perform differential expression analysis on samples that are normalized to ensure that expression values for a given gene are comparable across samples, and in particular to remove the potential effect of library size on the analysis. The null hypothesis of interest is that there is no mean difference between conditions on the data to be analyzed, i.e., the normalized data. Thus, these are the data that should be permuted, not the raw expression. Thus, the Li *et al* permutation scheme is not informative about the desired analysis on the normalized data. Our results indicates that the apparent false positives of `dearseq` using the Li *et al* scheme are likely detecting differences in library size. See Supplementary information for a detailed demonstration of the issues.

Both `limma-voom`[7] and `NOISeq`[8] also controlled FDR adequately using the amended permutation scheme (see Supplementary Figure S1) – note that this procedure is difficult for `voom-limma`, `edgeR`[2] and `DESeq2`[3] because normalization is baked into their analysis methodology. Supplementary Figure S2 shows that `dearseq` asymptotic test achieved higher power compared to both `limma-voom` and `NOISeq` (when *n >* 20 per group).

Furthermore, `dearseq` is capable of handling many experimental designs beyond the simple two conditions comparison setting of the Wilcoxon test, and thus constitutes a versatile option for differential expression analysis of large human population samples.

## Acknowledgements

Computer time for this study was provided by the computing facilities MCIA (Méesocentre de Calcul Intensif Aquitain) of the Universitée de Bordeaux and of the Universitée de Pau et des Pays de l’Adour.

## Availability of data and materials

All code and data needed to reproduce the results presented here are openly accessible from Zenodo with DOI 10.5281/zenodo.6536901 [10].

## Authors’ contributions

BPH and KB performed the numerical simulations and generated the figures. BPH, RT and DA designed the analysis and wrote the manuscript. All authors read and approved the final manuscript.

## Supplementary materials

### S1 Supplementary figures of performance evaluation

**Figure S1:**
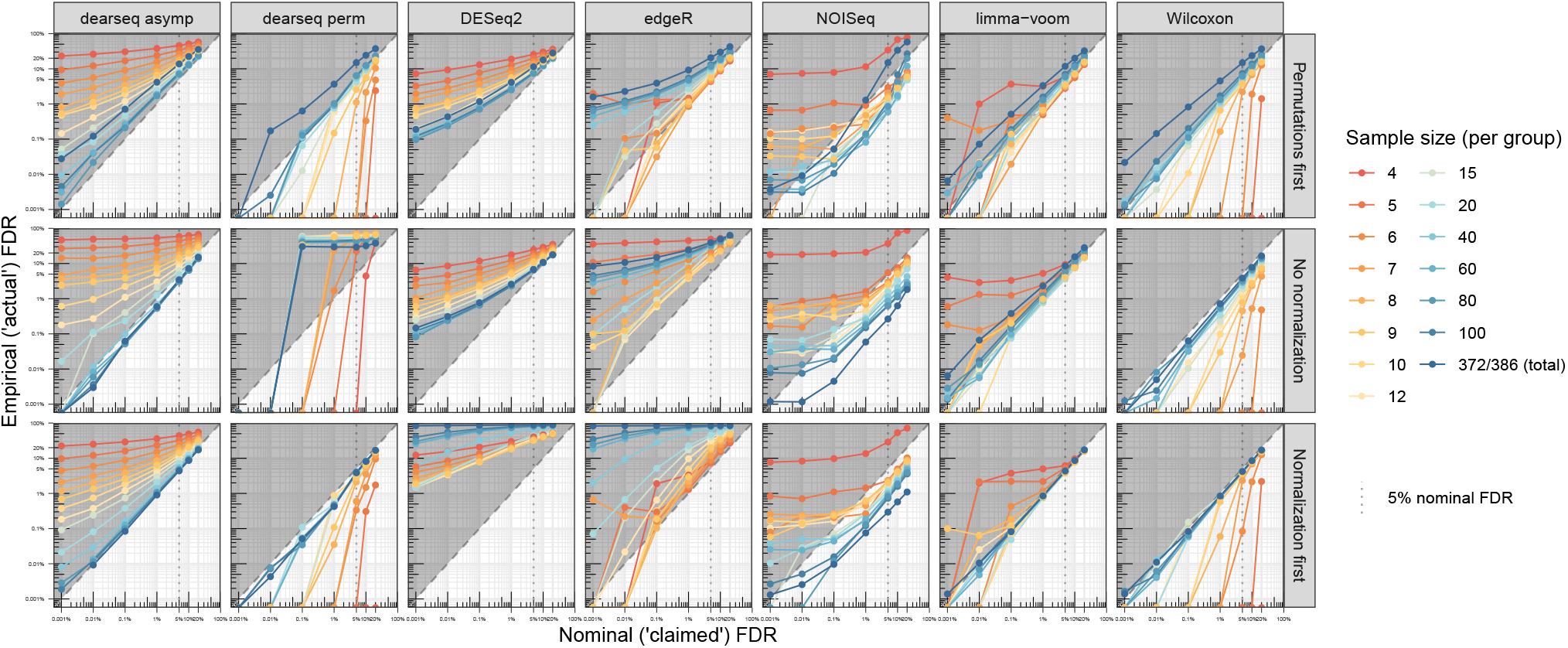
Empirical FDR control against Nominal FDR level by method,. Average over 50 semi-synthetic dataset generated from the *GTEx Heart atrial appendage VS Heart left ventricle* data. 50% of the true Differentially Expressed (DE) genes are randomly sampled in each semi-synthetic dataset (i.e. 2,889 genes remains unpermuted as true positives) and considered as gold-standard DE genes. **Panel A** reproduces the results from Li et al.[1] Figure 2A when all methods are applied to the same data (first permuted to generate null gene expression and then normalized) on the full sample size (372 and 386 samples in each group respectively). **Panel B** study the impact of both the sample size as well as the respective order between the data normalization and the random permutations to generate non-differentially expressed genes on the FDR control of the Wilcoxon test and on both asymptotic and permutation tests from `dearseq`.

**Figure S2:**
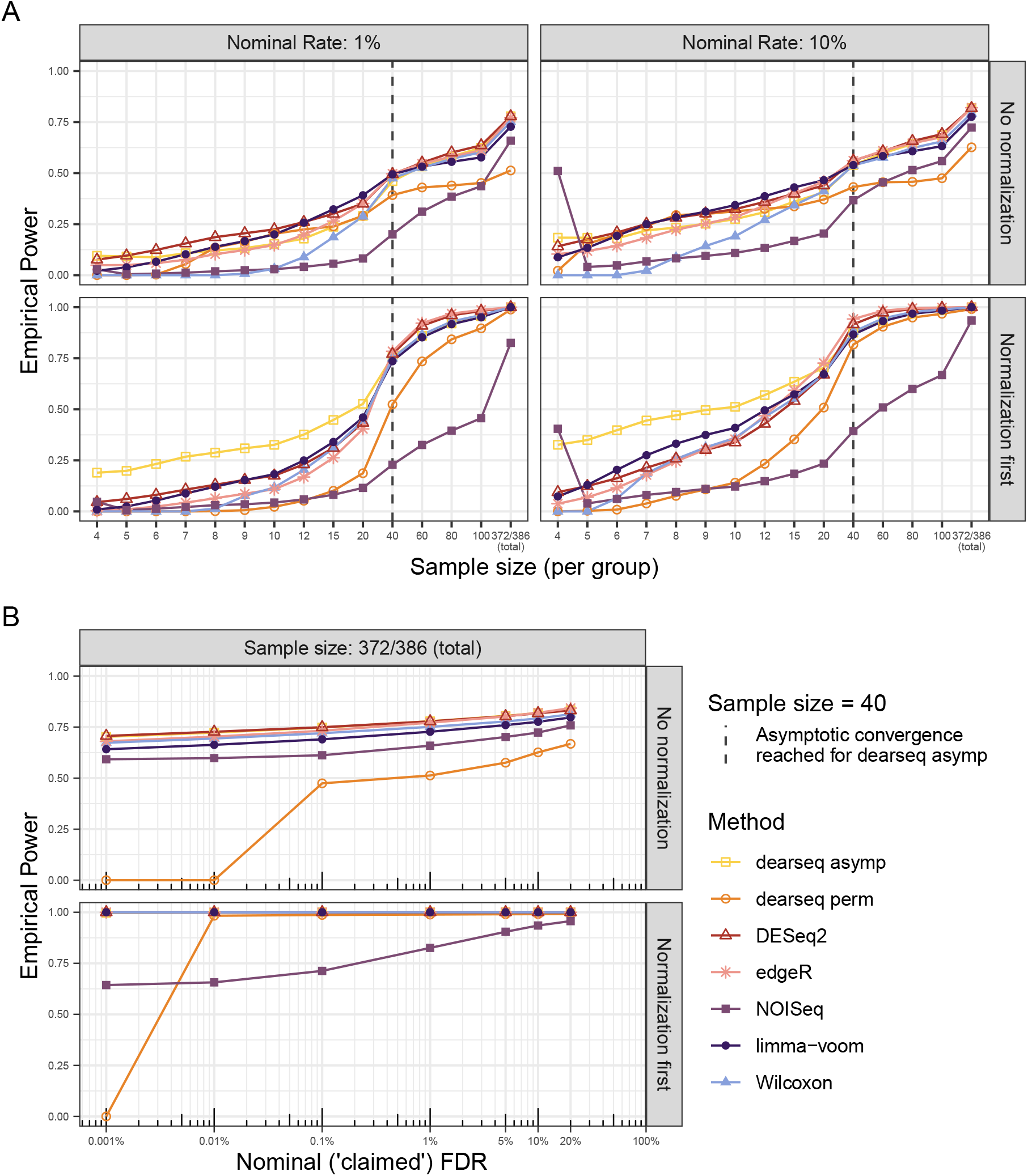
Empirical statistical power by method. Average over 50 semi-synthetic dataset generated from the *GTEx Heart atrial appendage VS Heart left ventricle* data. 50% of the true Differentially Expressed (DE) genes are randomly sampled in each semi-synthetic dataset (i.e. 2,889 genes remains unpermuted as true positives) and considered as gold-standard DE genes used as true positives. NB: for the empirical power to be interpretable, FDR control is warranted: thus **edgeR** and **DESeq2** results should not be interpreted here. **Panel A** reproduces the results from Li et al.[1] Figure 2B as a function of sample size for both 1% and 10% nominal FDR levels, when all methods are applied to the same data (either without any normalization or when the data are first normalized before randomly swapping values to generate expressions under *H*_0_). **Panel B** study the impact of the nominal FDR level in both cases for the full sample size (372 and 386 samples in each group respectively).

**Figure S3:**
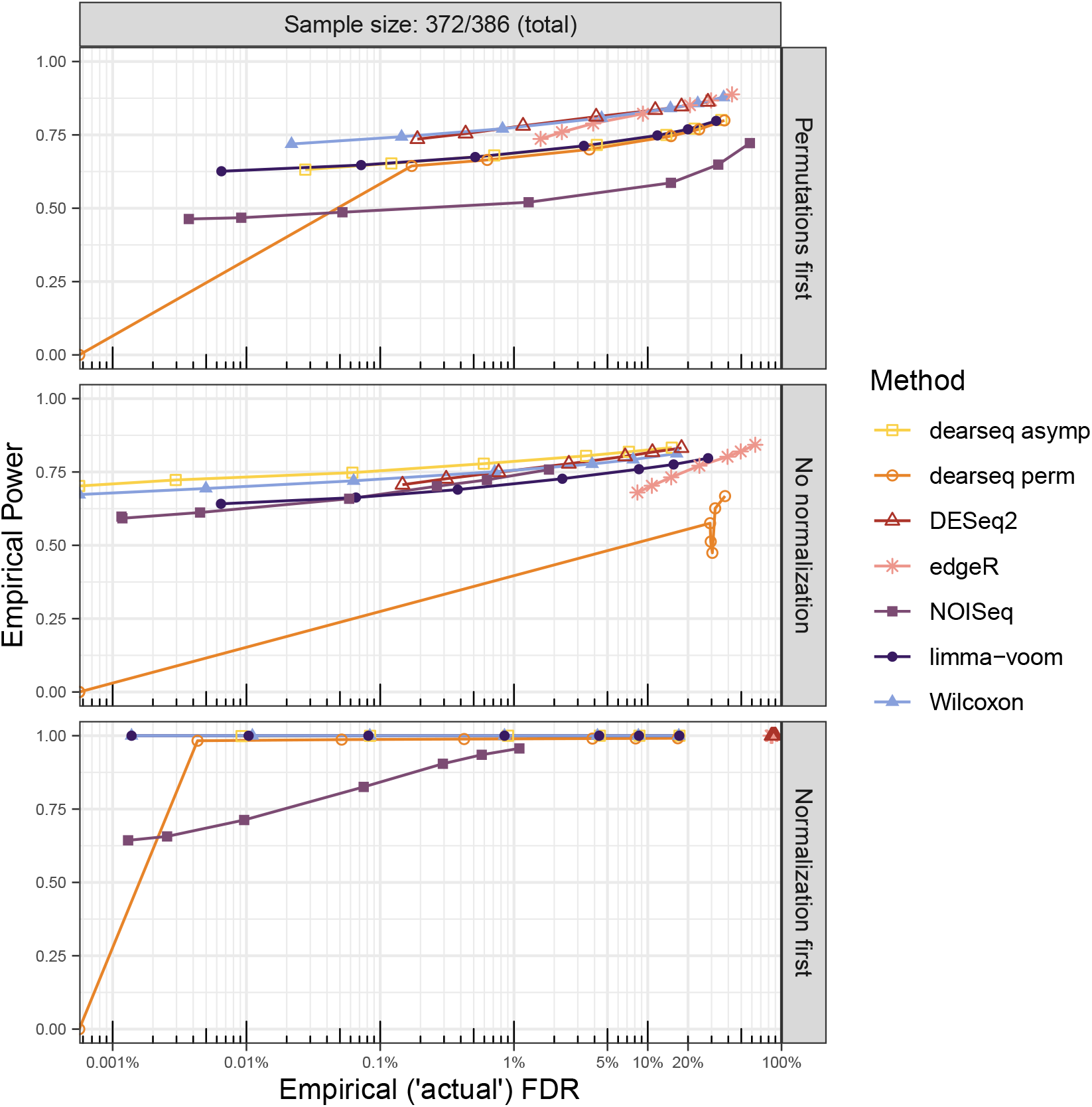
Empirical statistical power agianst empirical FDR by method. Average over 50 semi-synthetic dataset generated from the *GTEx Heart atrial appendage VS Heart left ventricle* data for the full sample size (372 and 386 samples in each group respectively). 50% of the true Differentially Expressed (DE) genes are randomly sampled in each semi-synthetic dataset (i.e. 2,889 genes remains unpermuted as true positives) and considered as gold-standard DE genes used as true positives.

### S2 Impact of library size and normalization

The main source of false positives generated in the *permutation first* scheme is likely the difference in library size. Contrary to when all genes are permuted (cf the analysis presented by Li *et al* [1] in their Figure 1 where neither `dearseq` nor `limma-voom` or `NOISeq` suffer from false positive inflation), when some genes – *a fortiori* differentially expressed (DE) genes – are left unpermuted, a difference in library size between the two condition of interest can subsist even after the permutation. In such case, this library size difference will affect the normalization.

Figure S4 displays and characterizes the imbalance of library sizes between the two heart conditions from the *GTEx Heart atrial appendage VS Heart left ventricle* data set used in this example. Figure S5 shows that this imbalance is mainly conserved in the subset of 5,778 genes that are considered as truly DE by Li *et al* (the intersections of genes that are significantly DE according to all five methods `DESeq2`, `edgeR`, `NOISeq`, `limma-voom` and Wilcoxon test at a FDR threshold of 10^−6^). This also explains results from their supplementary Figure S19 where the higher the proportion of true DE genes, the more false positives are generated by this library size difference remaining after their permutation scheme.

**Figure S4:**
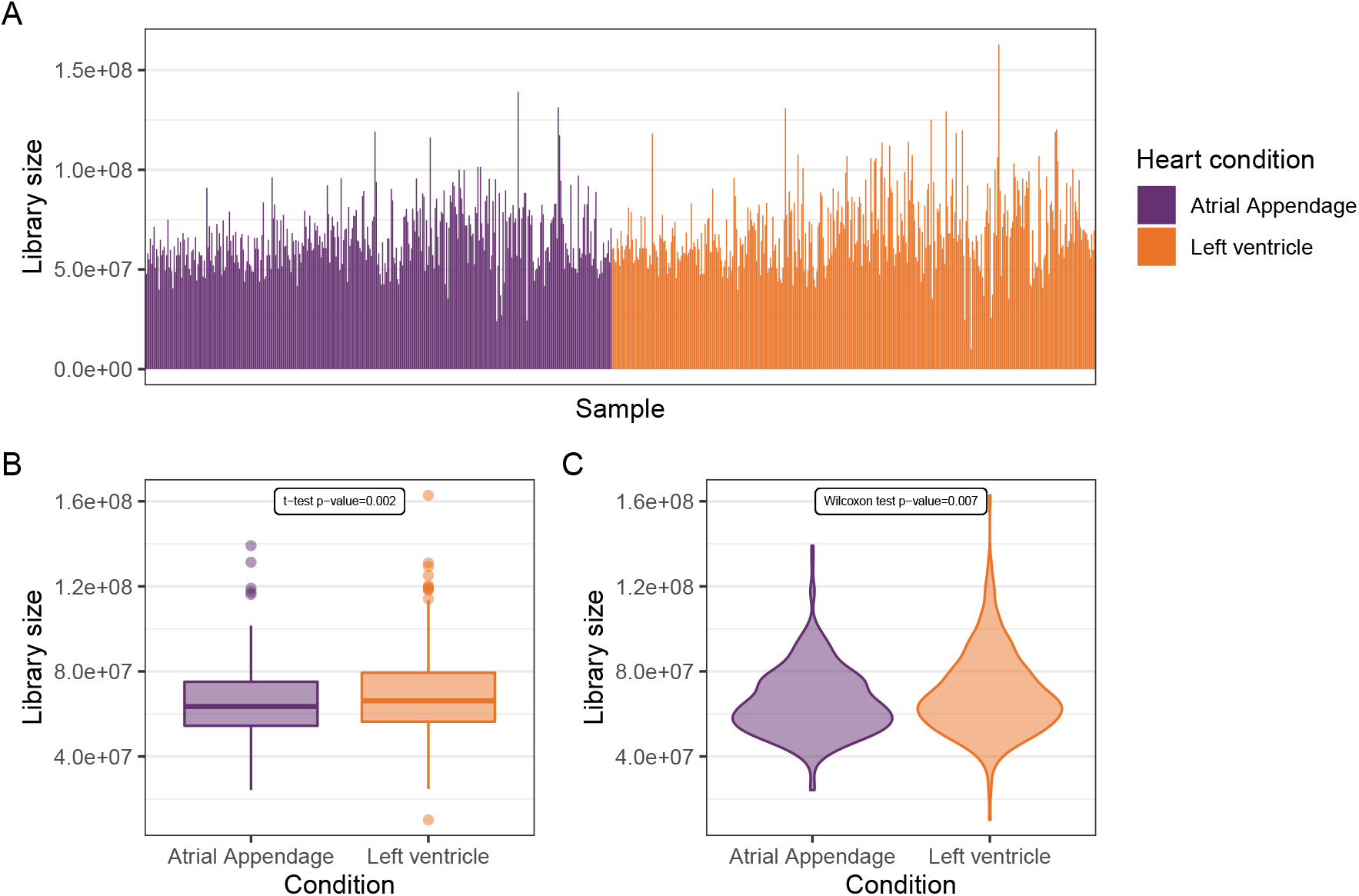
Library size differences in the *GTEx Heart atrial appendage VS Heart left ventricle* data. **Panel A** displays the library sizes of all 758 samples (372 and 386 in the atrial appendage and left ventricule heart conditions respectively). **Panel B** presents a boxplot highlighting the statistically significant difference with a t-test. **Panel C** presents a violin plot for a non-parametric comparison with the Wilcoxon test.

### Toy example

Here we present a toy example to illustrate the impact of post-permutation normalization due to library size imbalance.

**Figure S5:**
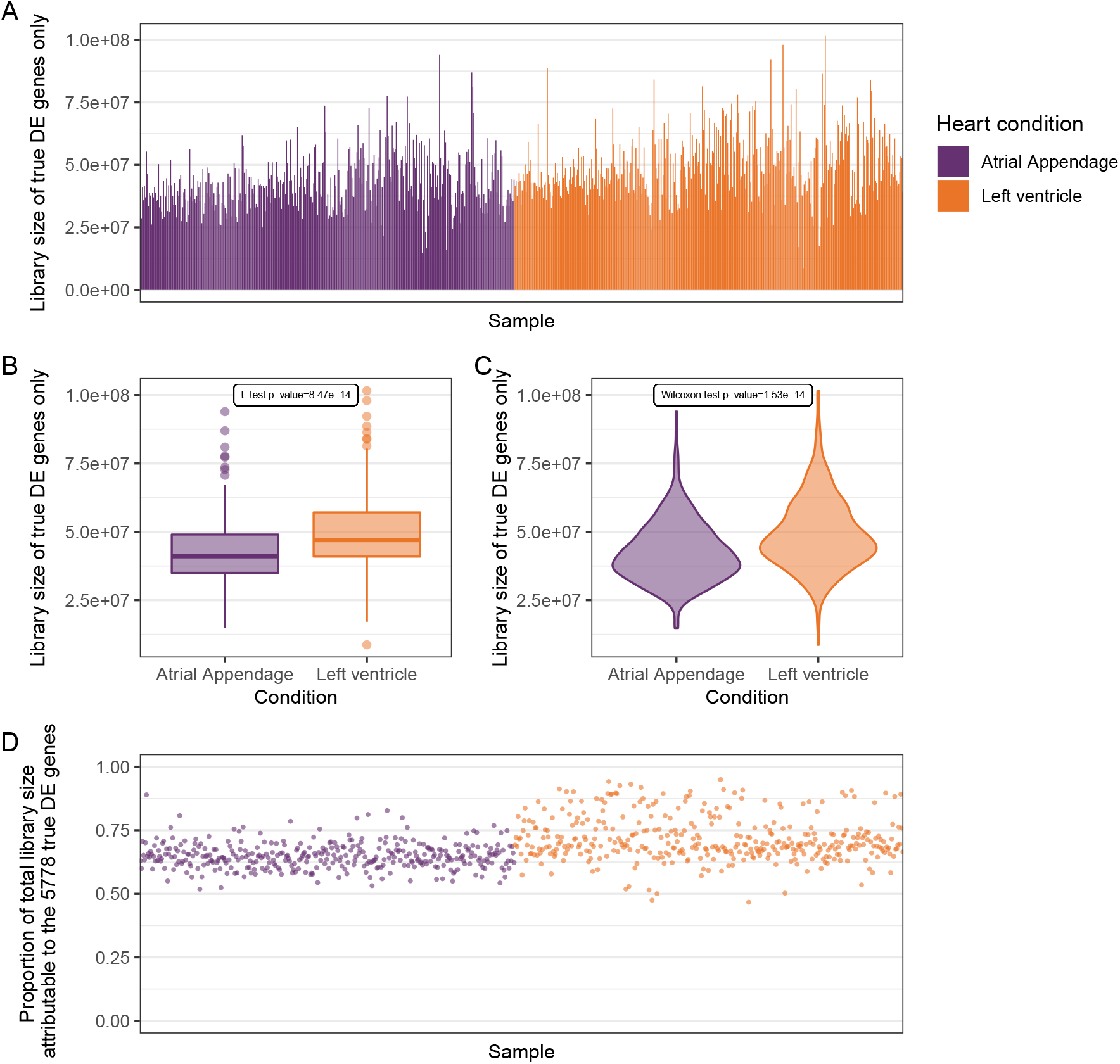
Library size of true DE genes only in the *GTEx Heart atrial appendage VS Heart left ventricle* data. **Panel A** displays the library sizes of all 758 samples (372 and 386 in the atrial appendage and left ventricule heart conditions respectively) when only using the 5,778 true DE genes. **Panel B** presents a boxplot highlighting the statistically significant difference with a t-test. **Panel C** presents a violin plot for a non-parametric comparison with the Wilcoxon test. **Panel D** shows that most of the total library size is accounted for by the subset of the 5,778 true DE genes, and even more so for the left ventricle heart condition

Consider three genes: *Gene 1* and *Gene 3* are truly associated with the condition of interest *A*, while *Gene 2* is independent of this condition. These three genes are measured across 10 samples with a library size that varies from 80 to 15,000. Table S1 presents the raw data while Tables S2, S3, and S4 display the normalized, permuted and normalized permuted data respectively. We observe that in this example there is confusion between the condition and the library size (i.e. not normalizing the data leads to *Gene 2* being significantly associated with condition *A*, while normalizing by 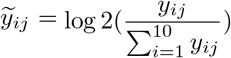 removes the spurious association). Following the Li *et al* [1] permutation scheme, we only permute *Gene 2*, and we keep *Gene 1* and *Gene 3* unchanged as both are true DE genes. While the permuted *Gene 2* is not associated to condition A (as expected), the permutation has swapped values coming from various library sizes: due to library size differences, expression values are nonexchangeable across samples under the null, as can be seen in Table S3. And because of *Gene 1* and *Gene 3* remaining unpermuted, the new library sizes are quite close to the original ones, still correlated to the condition of interest. Normalizing to account for library sizes that do not match with the swapped values in *Gene 2* now has the unwanted effect of adding back differences between the two conditions due to the confusion with the library size, as shown in Table S4.

**Table S1:**
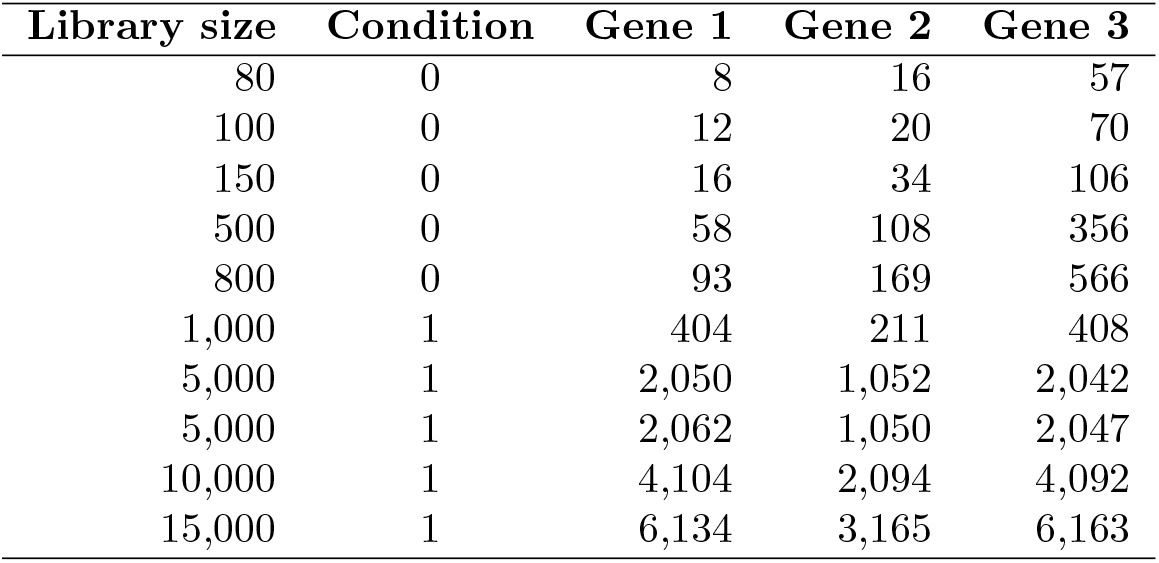
Original data. Wilcoxon test p-value for Gene 2 between the two conditions is 0.008.

**Table S2:** Normalized data. Wilcoxon test p-value for Gene 2 between the two conditions is 0.674.

| Library size | Condition | Gene 1 | Gene 2 | Gene 3 |
| --- | --- | --- | --- | --- |
| 80 | 0 | -3.32 | -2.32 | -0.49 |
| 100 | 0 | -3.06 | -2.32 | -0.51 |
| 150 | 0 | -3.23 | -2.14 | -0.50 |
| 500 | 0 | -3.11 | -2.21 | -0.49 |
| 800 | 0 | -3.10 | -2.24 | -0.50 |
| 1,000 | 1 | -1.31 | -2.24 | -1.29 |
| 5,000 | 1 | -1.29 | -2.25 | -1.29 |
| 5,000 | 1 | -1.28 | -2.25 | -1.29 |
| 1,0000 | 1 | -1.28 | -2.26 | -1.29 |
| 15,000 | 1 | -1.29 | -2.24 | -1.28 |

**Table S3:**
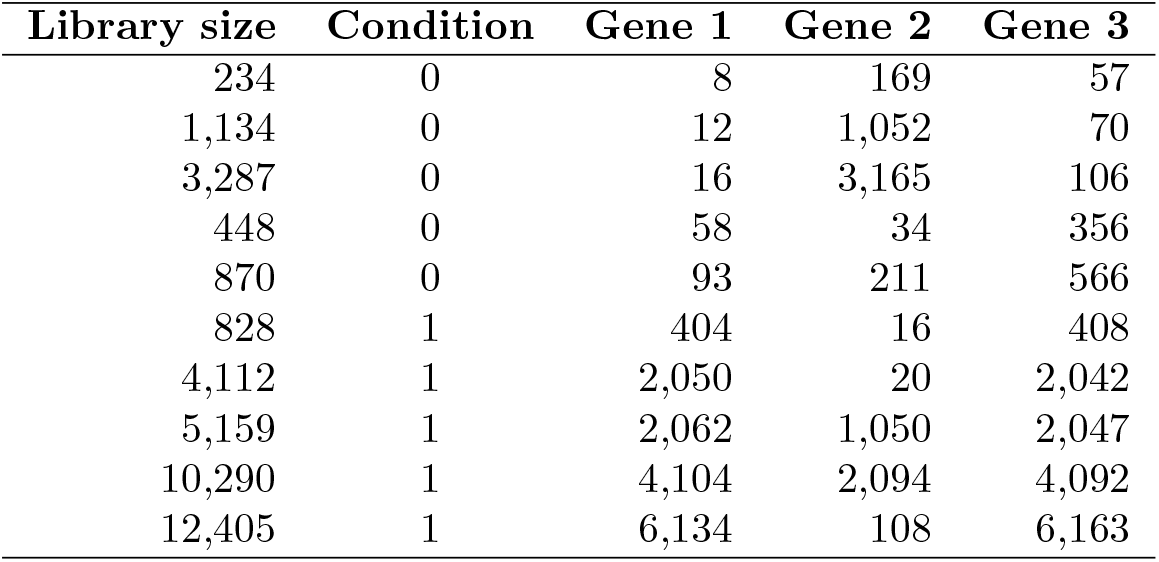
Permuted data. Wilcoxon test p-value for Gene 2 between the two conditions is 0.421.

| Library size | Condition | Gene 1 | Gene 2 | Gene 3 |
| --- | --- | --- | --- | --- |
| 234 | 0 | 8 | 169 | 57 |
| 1,134 | 0 | 12 | 1,052 | 70 |
| 3,287 | 0 | 16 | 3,165 | 106 |
| 448 | 0 | 58 | 34 | 356 |
| 870 | 0 | 93 | 211 | 566 |
| 828 | 1 | 404 | 16 | 408 |
| 4,112 | 1 | 2,050 | 20 | 2,042 |
| 5,159 | 1 | 2,062 | 1,050 | 2,047 |
| 10,290 | 1 | 4,104 | 2,094 | 4,092 |
| 12,405 | 1 | 6,134 | 108 | 6,163 |

**Table S4:** Normalized permuted data. Wilcoxon test p-value for Gene 2 between the two conditions is 0.031.

| Library size | Condition | Gene 1 | Gene 2 | Gene 3 |
| --- | --- | --- | --- | --- |
| 234 | 0 | -4.87 | -0.47 | -2.04 |
| 1134 | 0 | -6.56 | -0.11 | -4.02 |
| 3287 | 0 | -7.68 | -0.05 | -4.95 |
| 448 | 0 | -2.95 | -3.72 | -0.33 |
| 870 | 0 | -3.23 | -2.04 | -0.62 |
| 828 | 1 | -1.04 | -5.69 | -1.02 |
| 4112 | 1 | -1.00 | -7.68 | -1.01 |
| 5159 | 1 | -1.32 | -2.30 | -1.33 |
| 10290 | 1 | -1.33 | -2.30 | -1.33 |
| 12405 | 1 | -1.02 | -6.84 | -1.01 |

When repeating this toy example 500 times, the Wilcoxon test on normalized permuted data clearly has an inflated Type-I error as showed in Figure S6 compared to the uniform distribution expected under the null (p-values are uniform for the normalized data as well as for the permuted data, as expected).

**Figure S6:**
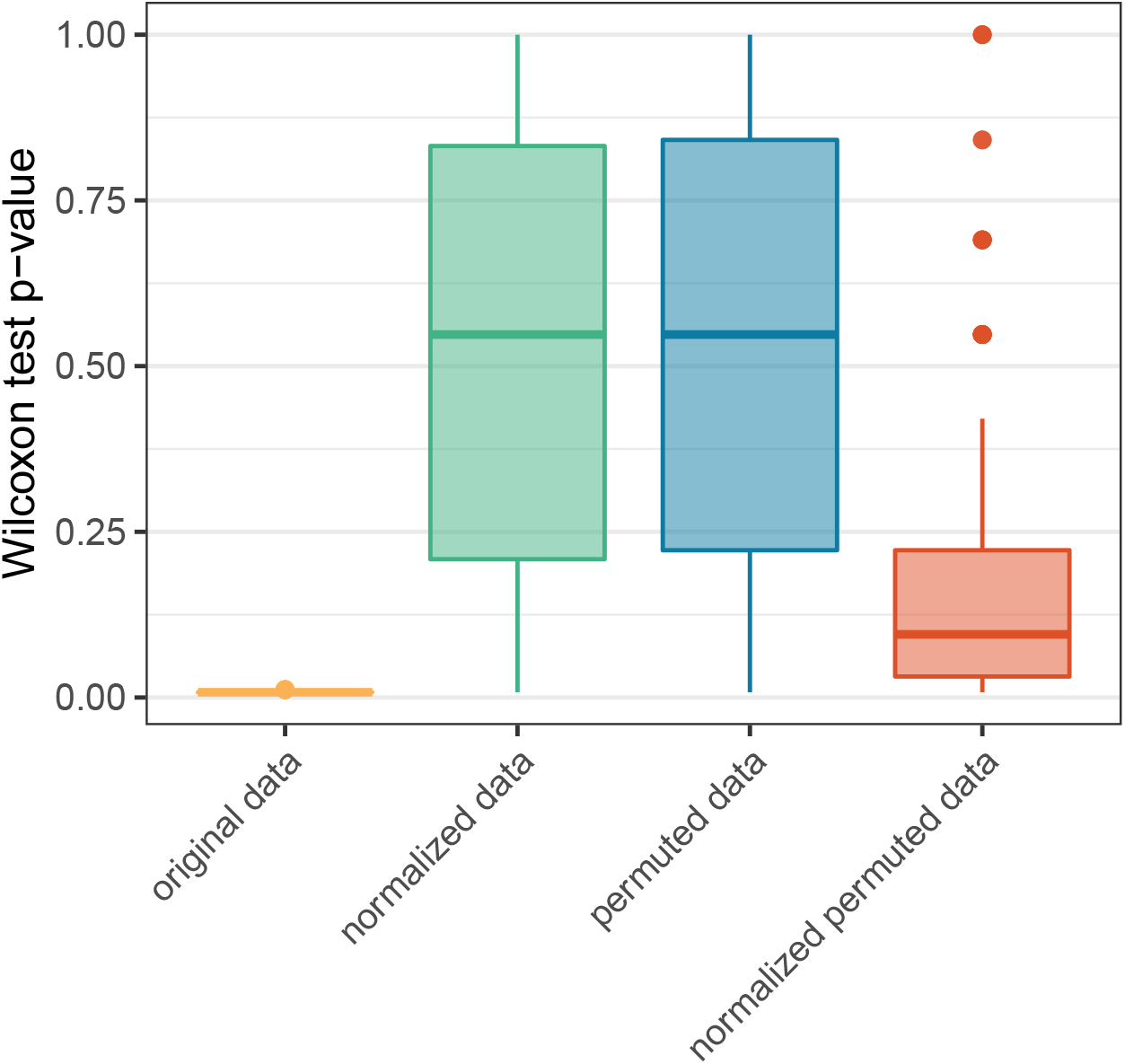
Boxplots of Wilcoxon test p-values according to the data processing in 500 repetitions of the toy example.

